# Dynamic whole-body models for infant metabolism

**DOI:** 10.1101/2024.11.25.625291

**Authors:** Elaine Zaunseder, Faiz Khan Mohammad, Ulrike Mütze, Stefan Kölker, Vincent Heuveline, Ines Thiele

## Abstract

Comprehensive, sex-specific whole-body models (WBMs) accounting for organ-specific metabolism have been developed to allow for the simulation of adult and infant metabolism. These WBMs are evaluated daily, giving insights into metabolic flux changes that occur in one day of an infant’s or adult’s life. However, for medical applications, such as in metabolic diseases and their treatment, an evaluation and concentration predictions on a shorter time scale would be beneficial. Therefore, we developed a dynamic infant-WBM that couples metabolite dynamics in short time frames through physiology-based pharmacokinetic models with the existing infant whole-body models. We then tailored the dynamic infant-WBM enabling the prediction of isovalerylcarnitine (C5), a clinical biomarker used for the inherited metabolic disease isovaleric aciduria (IVA). Our results show that, as expected, the predicted C5 concentrations exceeded the newborn screening thresholds during the time (36 - 72 hours) newborn screening blood samples are taken in the IVA models but not in models simulating healthy infants. We also demonstrate how the dynamic infant-WBMs can be used to test the effect changes in dietary intake have on the biomarker. Since the dynamic infant-WBMs were parametrised with literature-derived experimental or estimated values, we show how uncertainty quantification can be applied to quantify the parameter uncertainties. We found that the fractional unbound plasma needed to be estimated correctly, as this parameter strongly impacted C5 concentration predictions of the dynamic infant-WBMs. Overall, the dynamic infant-WBMs hold promise for personalised medicine, as it enables personalised biomarker concentration predictions of healthy and diseased infant metabolism in various time intervals.

## 2 Introduction

Newborn screening programs aim at the early detection of treatable rare diseases that pose a risk to the physical and mental development of affected children. Ideally, this identification should occur pre-symptomatically to significantly reduce morbidity and mortality, making these programs highly successful instruments of secondary prevention [1, 2, 3] Therefore, blood samples from newborns are collected on the first days of life (i.e., in Germany at 36 - 72 h of life) and sent to a newborn screening centre for analysis [4]. As soon as infants with an inherited metabolic disease (IMD) are identified, beneficial treatment can start. However, due to the substantial variability of IMDs, the disease management and therapy need to be adapted to the age-dependent requirements and the individual disease severity requiring carefully personalised treatments for each patient [5].

Numerous successive generic reconstructions of human metabolism have been published, such as the Recon series [6, 7, 8, 9] and the human metabolic reaction series [10], which presented the first generic reconstructions of the human metabolic network using cell-based models. Subsequently, sex-specific whole-body models of human metabolism (WBMs) have been constructed by integrating information from Recon3D, organ-specific details, and omics data [11]. These models capture the metabolism of 26 organs and six blood cell types in two, male and female, adult models named *Harvey* and *Harvetta* [11]. The goal of these WBMs is to model the metabolism of the entire human, and not solely on a cellular level, as seen in the Recon models [11]. All of these models focus on adult metabolism. To enable the investigation of infantile metabolism, the sex-specific infant whole-body models (infant-WBMs) have been developed [12] spanning the first 180 days of life. For the creation of these models detailed knowledge of infants’ physiology, metabolic processes, and newborn screening data was used. They account for organ-specific parameters and included in detail the energy demand for brain development, heart function, muscular activity, and thermoregulation. The models represent active infant metabolism and predict infant growth between zero and six months in agreement with growth trajectories by the WHO [13].

These generic, cell level, or whole-body genome-scale metabolic models can be used to predict reaction fluxes using constraint-based modelling [14], with flux balance analysis (FBA) being a preferred analysis method [15]. Therefore, steady-state is assumed [16], i.e., the concentration of metabolites in cells or organs remains constant over time, corresponding to an overall balance between metabolite inflow and outflow [15], and the simulation correspond to a specific time interval (e.g., daily). Consequently, this approach cannot be used to investigate short-term effects, such as responses to perturbations or dynamic variations in gene expression and metabolite levels. Such analysis would require the evaluation on shorter time scales with dynamic metabolite concentrations.

One possibility to capture the dynamic features and changes of human metabolism is through kinetic modelling, such as in physiological-based pharmacokinetic (PBPK) modelling [17, 18]. These PBPK models can be used to predict allosteric and post-translational regulation, alterations in metabolite concentrations, and considerations of thermodynamics [19]. PBPK models require extensive parametrisation, which often need to be estimated through extensive clinical studies. They describe absorption, distribution, metabolism, and excretion (ADME) of a particular drug or metabolite, while omitting other metabolic processes occurring in the human body. Consequently, combining dynamic and constraint-based models is of great interest as it would represent the dynamic nature of a particular pathway while accounting for the complexity of the overall metabolic network.

Combined dynamic and constraint-based models have been developed since they increase the spatial and temporal resolution of genome-scale metabolic models [19]. For instance, the unsteady-state FBA approach has been developed to improve the accurate prediction of metabolic flux states for red blood cells by relaxing the steady-state assumption [20]. Mannan et al [21] introduced an approach, which integrates parameters from a genome-scale metabolic network model into a kinetic model of the central carbon metabolism of *E. coli*. Another modelling framework considered both reaction kinetics and network connectivity constraints, emphasizing the role of metabolic network connectivity in influencing cellular control over metabolite levels [22]. For human metabolism, Mohammad et al [18] integrated PBPK modelling and constraint-based metabolic models to investigate the gut-brain axis for patients with an autism spectrum disorder. Moreover, Guebila et al [23] integrated a comprehensive PBPK model of glucose regulation by insulin, glucagon, and incretins with WBMs to perform dynamic flux balance analysis for type 1 diabetes. Furthermore, to analyse the effect of processes and genetic variations in ethanol metabolism, Zhu et al [24] integrated a PBPK model with the human whole-body models *Harvey* and *Harvetta* [11]. However, no dynamic whole-body model has been developed to analyse infant metabolism.

In medical applications, metabolic modelling is increasingly utilised, encompassing tasks, such as identifying drug targets [25] and off-target drug effects [9], studying cancer metabolism [26], enhancing the understanding of microbial interactions with the host organism [27], and also emerging as a field for investigating IMDs related to newborn screening [28]. IMD analysis is utilised in metabolic modelling to demonstrate the human metabolic model’s capability to accurately predict known biomarkers. Furthermore, a dietary treatment analysis with the infant-WBMs showed that these models can correctly simulate the effect of dietary interventions on patients with an IMD [12]. IMDs often involve a disruption of the normal metabolite flux [5]. Therefore, analysing the metabolic flux over time is significantly more informative for IMDs than the static measurement of intermediary metabolites [5]. For dynamic evaluations of IMD biomarker prediction in metabolic models, an approach combining PBPK modelling with cell-based metabolic human models has been developed [29].

Metabolic reconstructions try to mimic the dynamics in human metabolism as closely as possible but are limited by multiple heterogeneous sources of uncertainty [30]. Uncertainties in the metabolic reconstruction, such as the genome annotation, environment specification, biomass formulation, and network gap-filling, as well as the choice of flux simulation method can arise due to different choices and data used at these points [30]. To quantify these uncertainties, different approaches, including probabilistic metabolic reaction annotation [31], Monte-Carlo sampling [32, 33], and ensemble sampling [34], have been applied. Quantifying the uncertainty can test the reliability of the models along with the uncertainty associated with parameters and possible measurement errors coupled with their impact on the model’s output.

In this study, we developed a dynamic infant-WBM by coupling the metabolite dynamics from an infant-specific PBPK model, which simulates the change of biomarker concentration over time in different organs, the blood compartment, and the infant-WBM. This coupled model was then used to predict a known biomarker of an IMD over a specified time interval relevant for newborn screening. Moreover, we used the developed model to simulate the effect a reduction of a dietary amino acid intake has on the biomarker concentration. Further, we show how uncertainty quantification is used to analyse the reliability of the dynamic infant-WBMs and understand the impact of parameter uncertainties on the metabolite concentration prediction.

## 3 Results

### 3.1 Procedure to simulate dynamics with an infant-WBM-PBPK model

The development of dynamic infant-WBMs was based on the coupling of infant PBPK models [35] and sex-specific infant-WBMs [12] such that metabolite concentration predictions could be used to update the constraints on the fluxes within the infant-WBM model and vice versa (Figure 1). Therefore, we devised a four-step procedure to couple PBPK models and infant whole-body models such that dynamic evaluations of biomarkers and drugs are possible for individual infants (see Method section for details). This process first included the adaptation of an existing PBPK model [18] to represent infant physiology, including the appropriate parametrisation of organ and blood volumes, organ densities, the body surface area, height and blood flow rates [36, 37, 38]. The second step adapts compound-specific parameters, where the compound can either be a drug or a metabolite, e.g., a biomarker for an IMD. Therefore, parameter information, such as lipophilicity, molecular weight, and fractional unbound plasma rate, were used to calculate the compound-specific tissue partition coefficient [39, 40]. Further, it included information on urinary or faecal excretion of a compound [41]. The third step prepares for solving the system of ordinary differential equations (ODEs) of the PBPK model, which included defining parameters, such as frequency and time interval of the simulation. In the fourth step, the PBPK and WBM models were then iteratively coupled by i) setting the initial values for the compound concentration in each organ in the PBPK model, ii) computing the metabolite concentration with the PBPK model, iii) updating the corresponding bounds of the metabolite in each organ in the infant-WBMs with the predictions from the PBPK model, iv) optimising the updated infant-WBM for the desired objective function (e.g., metabolite accumulation in the blood), and v) using the predicted flux from the infant-WBM to update the PBPK model. From this point, step four is repeated until the end of the defined time interval is reached. For details on the developed process and the individual steps see the Method section and our code published at https://github.com/ThieleLab.

**Figure 1:**
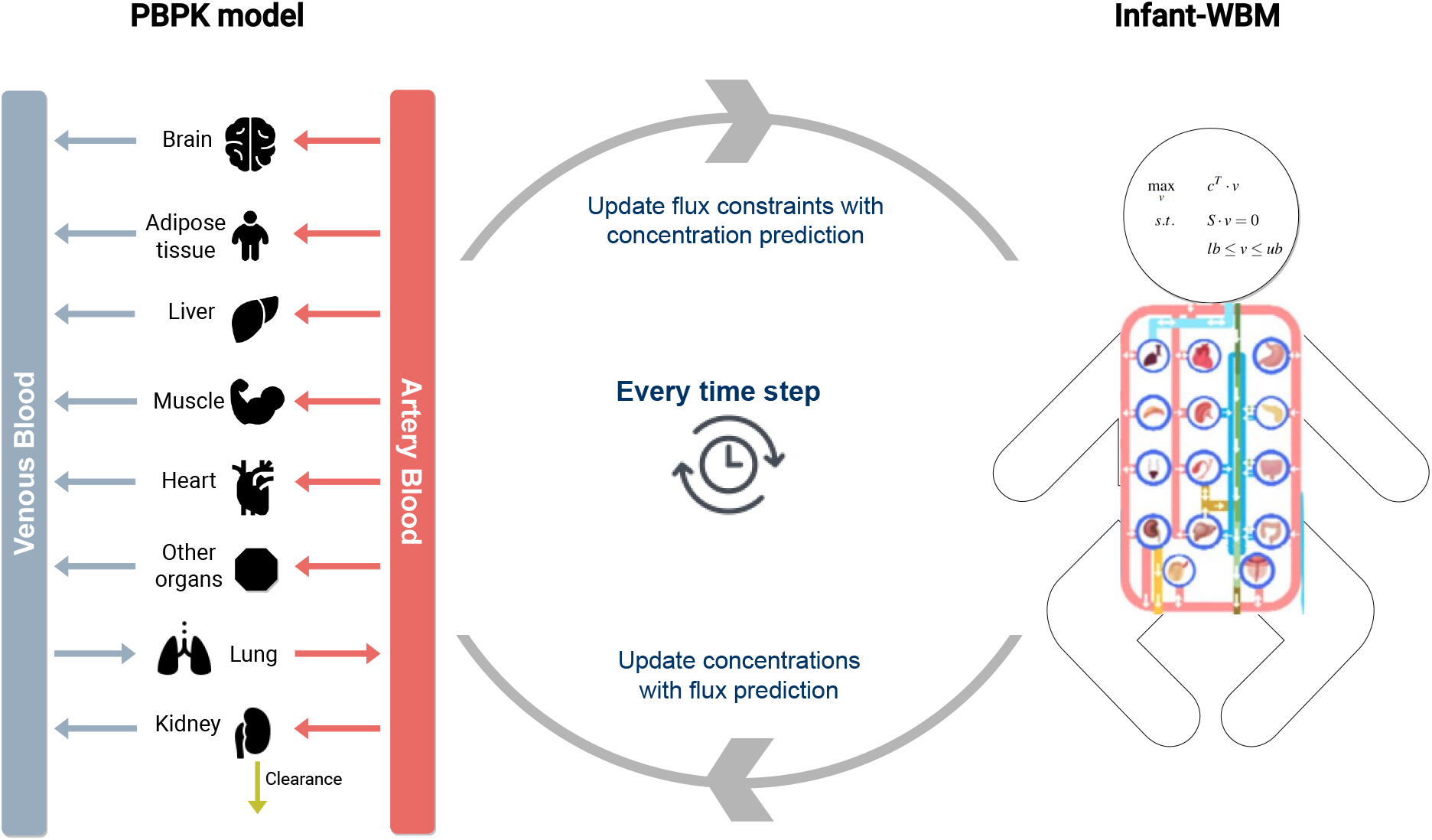
Schematic overview presenting the coupling of PBPK models [35] and infant-WBMs [12] to account for dynamic changes in metabolite concentrations.

### 3.2 Application of the procedure to dynamic infant-WBM for biomarker prediction

In the following, we demonstrate this four-step procedure by developing a dynamic infant-WBM for isovalerylcarnitine (C5), which is increased in the blood in patients with isovaleric aciduria (IVA, OMIM: #243500) [1, 42, 43]. Isovaleric aciduria (IVA) is caused by a deficiency in the activity of the mitochondrial FAD-dependent enzyme isovaleryl-CoA dehydrogenase due to biallelic pathogenic variants in the *IVD* gene (Entrez gene ID: 3712). The whole-body model includes all reactions necessary for the correct biochemical representation of IVA. In brief, isovaleryl-CoA (VMH ID: ivcoa) is generated in the mitochondrial compartment through the isovaleryl-CoA dehydrogenase reaction (VMH ID: ACOAD8m). Isovaleryl-CoA can then undergo further metabolism or be transported into the cytosol (VMH ID: IVCOAACBP), where it undergoes transesterification with carnitine, resulting in C5 (VMH ID: ivcrn) via the reaction (VMH ID: C50CPT1). C5 is then transported outside the cell (organ in our case, VMH ID: IVCRNe). Additionally, a further elimination route of isovaleryl-CoA exists via transesterification with glycine in the glycine N-acyltransferase reaction (VMH ID: RE2427M), yielding isovaleryl-glycine (VMH ID: CE4968). The virtual metabolic human identifiers (VMH IDs) referred to in this study represent the reaction names of the corresponding organ-specific reactions in the infant-WBM models. The non-organ-specific part of the VMH ID can be searched in the VMH database (https://www.vmh.life) for further information.

#### Step 1: Adaptation of an existing PBPK model for C5 prediction

First, a published PBPK model [44] was obtained which describes the dynamics of a system with 23 ODEs. These ODEs describe the time-dependent compound concentration in 14 tissues (such as adipose tissue, brain, small intestine, large intestine, heart, kidney, liver, lung, muscle, pancreas, skin, spleen, stomach, and bone), two blood compartments (artery and venous blood), two excretion compartments (renal and bile), as well as the transit from bile to small and large intestine lumen (see Method section for further details). We extended the PBPK model for the metabolic activity of the isovaleryl-CoA dehydrogenase reaction by assuming Michaelis-Menten kinetics. Therefore, we retrieved experimental data from the literature (*K*_*m*_ = 125 µmol [45], Table S2 (D)). The maximum velocity of the compound-producing reaction was fixed to *V*_*max*_ = 0.225 mol/min.

To allow for the iterative coupling between the PBPK and the infant-WBM (in Step 4), the overlapping organ and reactions in the PBPK and the infant-WBM models were identified. In the infant-WBM, exchange reactions (VMH ID: EX ivcrn(e) [bc] between organs and the blood compartment existed for C5 (VMH ID: ivcrn) in the lung (Lung EX ivcrn(e) [bc]), heart (Heart EX ivcrn(e) [bc]), muscle (Muscle EX ivcrn(e) [bc]), and kidney (Kidney EX ivcrn(e) [bc]). The PBPK model was reduced to these organs, the blood compartment, and the respective excretion pathways, resulting in the following reduced PBPK model for dynamic C5 prediction. The isovaleryl-CoA dehydrogenase reaction (VMH ID: ACOAD8m) was present in these four organs, as well as 15 other organs in the infant-WBMs. Hence, the flux through this reaction was coupled for the overlapping organs between the reduced PBPK model and the infant-WBMs. At the end, the PBPK part of the coupled model consisted of the following mass-balance equations:

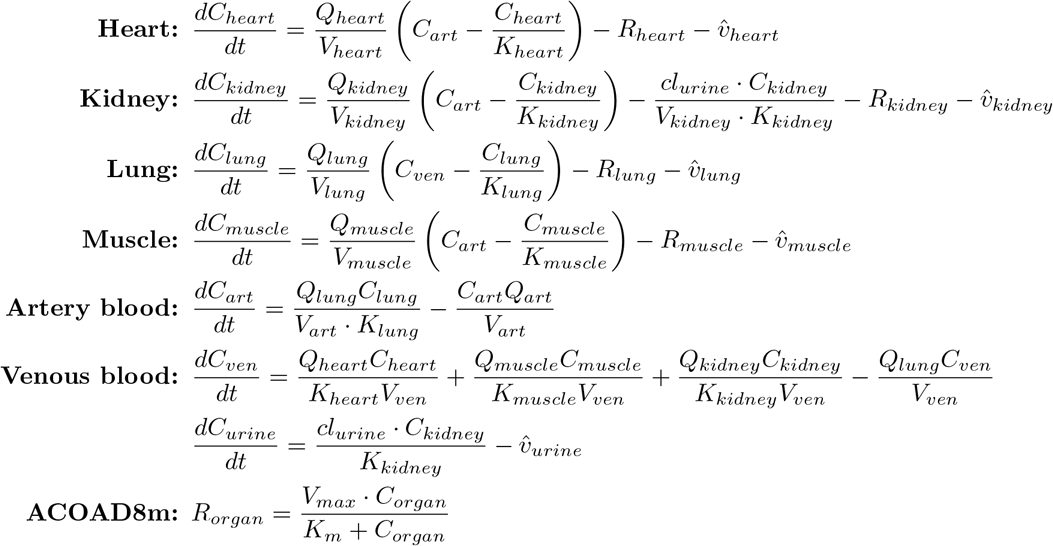

where *C*_*T*_ represents the concentration of C5 in a specific tissue T, *V*_*T*_ is the tissue volume, *Q*_*T*_ the blood flow rate, and *K*_*T*_ the calculated organ-specific tissue partition coefficient for C5. Furthermore, *cl*_*urine*_ represent renal clearance for C5. 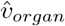 is the predicted flux value for each C5 organ-specific exchange reaction in the infant-WBM for each organ (see Method section for more details).

Next, we adapted this PBPK model for C5 by adjusting individual-specific, physiological parameters. To acquire these parameters for infants, we obtained the female and male infant-WBMs at age day one from the infant-WBM resource [12]. The PBPK model was assigned a body weight of 3.3 kg for male infants and 3.2 for female infants. Based on the body weight and measured references from newborns [46], the organ weights and organ-specific blood flow rates were determined as follows. The organ flow rates were obtained from the organ flow rate parameters used for the infant-WBM (SI material in [12]), which were reported in l/min/kg tissue and, hence, needed to be converted into l/h/organ [12] (Table S2 (A), (B)). The blood flow rate for venous and arterial blood was estimated as 3 l/h, since the blood flow rate ranges from 10 to 50 ml/min in infants weighing less than 5 kg [38], leading to an estimation of 0.6 - 3 l/h by the multiplication with 60/1000. The calculation of the organ volumes was based on the organ weights reported in the infant-WBMs and the respective organ density was obtained from literature [36] (Table S2 (C)). The venous and arterial blood volume estimations were based on the total blood volume resulting in 144 ml of blood in both compartments for a 3.3 kg newborn at day 1 (total blood volume: 288 ml). This value was comparable to the blood range of newborns at 72 hours of age (75 to 107 ml/kg) ranging between 248 - 353 ml for a 3.3 kg infant [47]. The body surface area was estimated as 0.2141*m*^2^ using the Mosteller method [48], with the average height of 50 cm for male newborns [37]. Similar to Toroghi et al [29], the physiological and physicochemical properties in the PBPK model were assumed to be constant over time.

#### Step 2: Determination of compound-specific parameters for C5 prediction

Next, we identified the C5-specific parameters from the literature and databases, such as the Human Metabolome DataBase (HMDB) [40] (Table S2 (D). The molecular weight of C5 is 245.3153 g/mol [40]. Further, a predicted lipophilicity value *l* = *−*2 was obtained from the HMDB [40]. The fractional unbound plasma was determined in experimental studies as 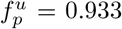 for all tissues *T* [49]. Similar to the study from Toroghi et al [29], the blood-to-plasma ratio was set to *B* = 1.

Based on these parameters and predefined specific volume fractions of neutral lipid, phospholipid, and water in tissue and plasma [50], the tissue partition coefficient *K*_*T*_ was calculated for C5 for the different tissues *T* using the Poulin method [39] (Table S2 (E)). Furthermore, we assumed that C5 was only excreted through urine, and not through faeces, since HMDB only listed urine as excreta in the disposition section for C5 (HMDB0000688) [40]. The renal clearance of acylcarnitines, such as C5, is 4 - 8 times higher than the one of C0 [51], which is reported to be 1-3 ml/min per adult person [41]. Taking 2 ml/min and dividing this by the average adult body surface area of 1.73*m*^2^, the infantile renal clearance of C0 would be 0.67 ml/min. Hence, the final infantile C5 renal clearance was set to be 4.64 ml/min, which is six times higher than the renal clearance of C0.

#### Step 3: Defining frequency and time intervals for C5 prediction

We defined that we would like to simulate the PBPK model over, e.g., two-hours, with one simulation corresponding to 10 minutes, during which the flux through 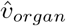 was assumed to be constant.

#### Step 4: Coupling of PBPK and infant-WBM models for C5 prediction

The setup from Step 3 means that after after ten minutes, the venous and arterial C5 concentration predictions from the PBPK model was used to update the bounds of the corresponding reactions in the infant-WBM (see Step 1 and Method section). Then, the infant-WBM was solved using flux balance analysis (FBA) using the COBRA Toolbox [16] in MATLAB, while maximising for the whole-body biomass reaction. From the resulting flux vector, we retrieved the flux value for each C5 organ-specific exchange reaction (Step 1, Methods) leading to a 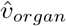 for each organ, which were used to update the PBPK model. Then, the ODE’s were solved in MATLAB. This step was repeated 12 times for the two-hour interval, 30 times for the five-hour interval, and 432 times for the 72-hour interval. The initial concentration of C5 was set to 0.1 µmol/l, the healthy average of over 2 million newborns [52]. This initial concentration was set for all tissues *T*.

### 3.3 Dynamic biomarker prediction for infant models

#### 3.3.1 Dynamic changes of C5 in biofluids and organs

To investigate whether the C5-specific dynamic infant-WBMs could be used to predict the time-dependent changes of C5 in the different biofluids and organs, we used the healthy female and male infant models and predicted the C5 concentration in the blood compartment. The predictions over a two-hour interval for the artery, heart, kidney, lung, muscle, and venous blood compartment showed similar behaviours for the organs in the female and the male models (Figure 2 (A, B)). Overall, the C5 concentrations were dynamically increasing or decreasing in the first 30 minutes until they were nearly constant, i.e., they reached a steady state. The female model obtained higher concentration values for the venous (0.38 µmol/l) and lung (0.33 µmol/l) than the male model (venous: 0.31 µmol/l; lung: 0.19 µmol/l) after the two-hour interval (Figure 2 (A, B)). For both models, the C5 concentration in the artery, heart, kidney, and muscle were lower than the initial value 0.1 µmol/l (Figure 2 (A, B)). All of the predicted values in the male and female models were beneath the 99^*th*^ percentile (0.51 µmol/l) of blood C5 concentration used as cut-off values for IVA [53, 43].

**Figure 2:**
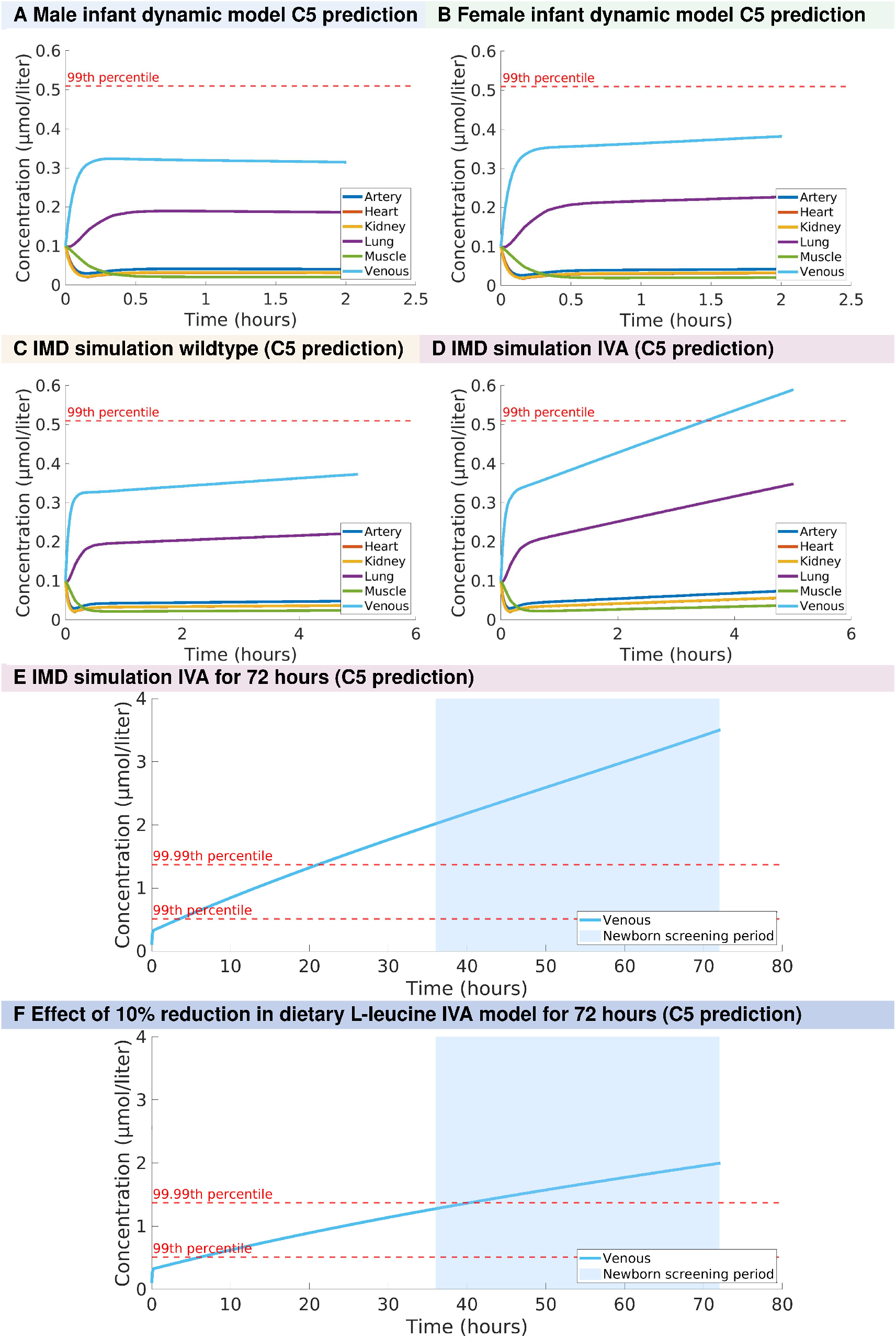
Concentration prediction of C5 from the dynamic infant-WBM for the artery and venous blood compartment, as well as the heart, kidney, lung, muscle over different time intervals. For (A) a male dynamic infant model and (B) a female dynamic infant model both optimising the growth rate over a two-hour interval. For an IMD analysis comparing (C) a wild type (WT) model and (D) an IVA model. And (E) an IVA model over a 72-hour interval also displaying the time interval, during which newborn screening is usually performed. (F) Predicted changes of C5 in IVA model over a 72-hour interval upon 10% dietary leucine reduction. Note that no further changes have been made to the IVA model. The dashed lines show the 99^*th*^ percentile (0.51 µmol/l) and 99.99^*th*^ percentile (1.37 µmol/l) of blood C5 concentration used as cut-off values for IVA [53]. 7

#### 3.3.2 Simulation of IVA and the predicted C5 concentration

To evaluate the consequences of a deficiency in the isovaleryl-CoA dehydrogenase, we derived an IVA model from the healthy model, by setting the bounds of the corresponding reactions in any organ to zero (Table S3, see Methods), thereby assuming a complete knockout. In the IMD simulation for IVA, we maximised for C5 accumulation in the blood compartment (i.e., the DM ivcrn[bc] reaction) in both, the healthy and the IVA model. Over the five hours of prediction time, the healthy (WT) model reached a C5 level of 0.37 µmol/l in the venous compartment (Figure 2 (C)). In contrast, in the IVA model, the C5 predictions in all organs increased continuously with the venous blood compartment having the highest concentration, reaching a level of 0.59 µmol/l after five hours (Figure 2 (D)). This value is higher than the 99^*th*^ percentile (0.51 µmol/l), which has been established in a large-cohort of healthy newborns to be a suitable cutoff for (attenuated) IVA [53]. Furthermore, the predicted concentration in all other organs was higher in the IVA than in the healthy model, lung (WT: 0.22 µmol/l, IVA: 0.35 µmol/l), heart (WT: 0.04 µmol/l, IVA: 0.06 µmol/l), kidney (WT: 0.04 µmol/, IVA: 0.06 µmol/l, and muscle (WT: 0.02 µmol/l, IVA: 0.04 µmol/l, Table S1). When predicting C5 concentration in the venous blood compartment over 72 hours, it reached 3.5 µmol/l (Figure 2 (E)) and thus exceeded the 99.99^*th*^ percentile (1.37 µmol/l) of blood C5 concentration, which had been established as a suitable cut-off value for IVA [53]. This cutoff value was exceeded after 22 hours and is thus elevated during the recommended age for newborn screening in Germany of 36 - 72 hours of life [54].

#### 3.3.3 Effect of reduced dietary L-leucine on C5 concentration in IVA models

The treatment of IMDs, such as IVA, needs to be individualised depending on age-dependent requirements and disease severity for each patient due to the substantial variability of IMDs [5]. These treatments include dietary interventions, such as restricting the uptake of metabolites or giving supplements [55].

Identifying the best treatment strategy for a patient, including the amount of natural food and supplements, is part of the patient-specific diet [56]. Hence, the possibility to test and evaluate therapies and treatments *in silico* with personalised models could be very beneficial for both patients and clinicians. As a first step to show the impact of dietary interventions on dynamic infant-WBMs, we investigate whether the predicted changes in C5 from the IMD analysis for IVA can be attenuated by dietary changes. For IVA, a study investigating 140 patients with IVA from 39 centres reported that 133 patients (from 38 centres) were given a protein restricted diet supplemented with leucine-free amino acids [57]. To analyse the effect a reduced dietary L-leucine uptake has on the dynamic model, we reduced the dietary intake of L-leucine in the male infant IVA model by 10% and predicted the C5 concentration over 72 hours. Note that we did not consider any further diet modification, such as carnitine supplementation, which is routinely done [57]. As expected, the reduction in dietary L-leucine led to a decrease in C5 venous blood concentration over the simulated time period of 72 hours (Figure 2 (F)). After 72 hours, the C5 venous concentration only increased to a value of 2 µmol/l, whereas the untreated dynamic IVA infant-WBM predicted a C5 venous concentration of 3.5 µmol/l (Figure 2 (E)). However, the C5 concentration remained still very high compared to the healthy C5 concentration. Taken together, this example demonstrates that the dynamic infant-WBM can be used for testing the effect of a single dietary change on a biomarker but also that further model refinement is required to reproduce known biomarker dynamics quantitatively.

### 3.4 Uncertainty quantification in dynamic infant-WBMs

The infant-specific PBPK model of the dynamic infant-WBMs was parametrised with values obtained from the literature, which could introduce uncertainties in the model. Therefore, we performed an uncertainty quantification (UQ) analysis for specific parameters using a Monte Carlo sampling method in the male infant-specific PBPK model [58]. For the UQ, we only analysed the infant-specific PBPK part of the coupled dynamic infant model, since this made it computationally possible to simulate 100,000 realisations in less than 15 minutes by using the Parallel Computing Toolbox implemented in MATLAB [59]. By quantifying this uncertainty and its impact on the model’s result, the reliability of the model when confronted with real-world data errors can be estimated. We investigated the following literature-derived parameters: lipophilicity *l*, urinary clearance *cl*_*urine*_, and fractional unbound plasma 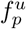. Therefore, we applied a Monte Carlo simulation with 100,000 realisations for every parameter and compared 100 randomly selected realisations with line plots to assess the impact of each uncertain parameters over a two-hour interval (Figure 3). In the previous section, the lipophilicity was assumed to be -2 [40]. The fractional unbound plasma was determined based on experimental studies performed in the literature as 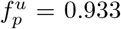 for all tissues *T* [49]. The renal clearance of C5 was estimated as 4.64 [51, 41]. With these parameters fixed, the infant-specific PBPK model predicted a C5 concentration in the venous compartment of 0.356 µmol/l after two hours. For each Monte Carlo evaluation, the parameters that were not randomly sampled were assigned these values.

**Figure 3:**
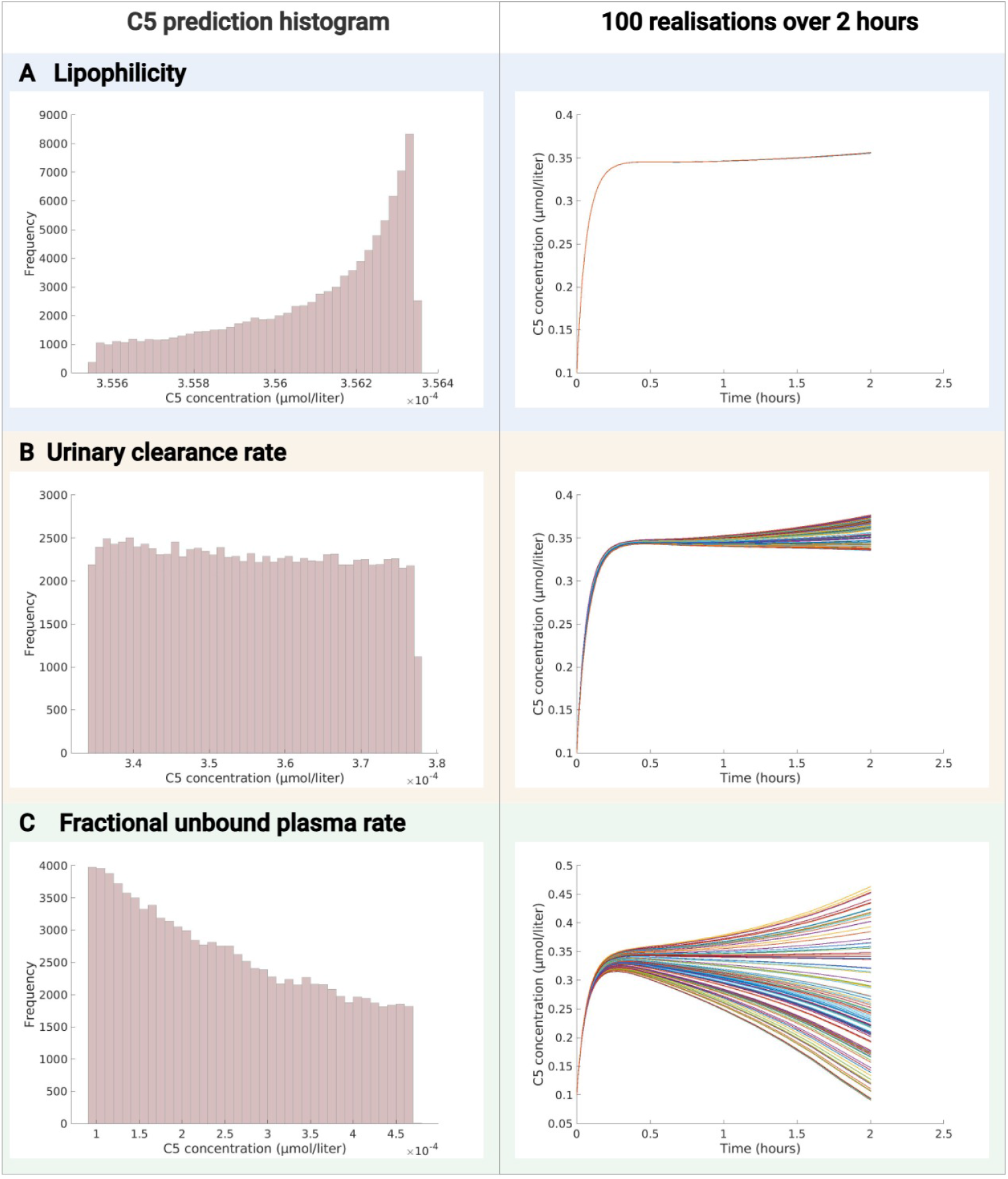
Histograms of 100,000 random initialisation of (A) lipophilicity parameter, (B) urinary clearance rate, and (C) fractional unbound plasma evaluated in the venous blood concentration prediction after 2 hours and 100 realisations of two-hour C5 predictions with randomly initialised respective parameter.

#### 3.4.1 Lipophilicity

The lipophilicity parameter refers to the capacity of C5 to dissolve in lipids [50]. The lipophilicity has been predicted as *l* = −2 using ALOGPS (https://vcclab.org/lab/alogps/) and *l* = −3 using ChemAxom (https://chemaxon.com/calculators-and-predictors#logp_logd) according to the HMDB [40] (https://hmdb.ca/metabolites/HMDB0000688). We applied uniform sampling in the interval *I*_*l*_ = [−3, −2] to obtain 100,000 random samples for the lipophilicity parameter 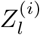, *i* = 1, …, 100, 000. These samples were then used as lipophilicity input parameters for the PBPK model to predict the C5 concentration over two hours in the venous compartment. The UQ analysis showed that the mean of the predicted venous concentration was 0.356 µmol/l (range: 0.355 - 0.356 µmol/l, Figure 3 (A)). Although the underlying distribution of the random variables 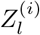 was uniform, the C5 prediction was negatively skewed, *s* = −0.77, (calculated with *skewness*.*m* [59], Figure 3 (A)). Overall, the C5 predictions over two hours of 100 realisations showed minimal variability in the C5 prediction over time due to the variation in the lipophilicity parameter *l* (Figure 3 (A)).

#### 3.4.2 Urine clearance rate

The urine clearance rate *cl*_*urine*_ describes the amount of urine excreted by an infant per time unit. The urine clearance rate of C5 was 4-8 times higher than the renal clearance of C0 [51], estimated as 1.16 ml/min/1.73m^2^ [41]. Consequently, the interval boundaries for the renal clearance were between 4.64 and 9.28, rounded to the interval *I*_*cl*_ = [4.5, 9.5]. Uniform sampling in this interval was applied to obtain 100,000 random samples for the renal clearance parameter, which was multiplied by the infant-specific body surface area. The resulting value was used for the renal clearance parameter 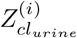, *i* = 1, …, 100, 000 to predict the C5 concentration over two hours in the venous compartment. The mean of the predicted concentration was 0.35 µmol/l, and the values were between 0.34 - 0.38 µmol/l (Figure 3 (B)) with a nearly uniform distribution that had a slight positive skewness, *s* = 0.04. The C5 predictions over two hours of 100 realisations showed that there was some variability in the C5 prediction over time due to the variation in the urine clearance rate *cl*_*urine*_ (Figure 3 (B)). For the first 30 minutes, the predictions were very similar for all 100 realisations, and with progressing time, the predictions showed more variability (Figure 3 (B)).

#### 3.4.3 Fractional unbound plasma

The fractional unbound plasma, 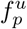, defines a drug’s binding degree in plasma [60]. It was determined based on a standard addition experiment and was reported to bess 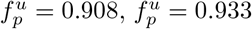, and 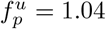 depending on the amount of added compound [49]. Therefore, uniform sampling was applied in the interval *I*_*f*_ = [0.9, 1.05] for 100,000 samples, 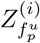, *i* = 1, …, 100, 000. These samples were then used as fractional unbound plasma input parameters for the PBPK model to predict the C5 concentration over two hours in the venous compartment. The mean of the predicted concentration was 0.25 µmol/l and the values were between 0.09 - 0.47 µmol/l (Figure 3 (C)). Unlike the underlying uniform distribution of the random variables 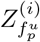, the model output 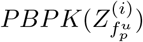 showed a positive skewness *s* = 0.3. Already starting after 10 minutes of prediction time, the 100 evaluated realisations showed variation in the concentration prediction, which increased with progressing time (Figure 3 (C)).

#### 3.4.4 Comparison of all uncertain parameters

All three investigated parameters, lipophilicity *l*, urinary clearance *cl*_*urine*_, and fractional unbound plasma 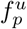, had uncertainties associated with them based on the methods and literature sources used to determine them. The UQ evaluation showed that the model’s C5 prediction varied in the venous blood compartment, but also in other compartments, depending on the investigated parameter (Figure 4 (A)). The variation in the venous C5 prediction due to the random sampling of the lipophilicity parameter *l* and urine clearance rate *cl*_*urine*_ was small compared to the output variation, which was due to the fractional unbound plasma 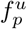 (Figure 4 (A)). The expected mean value of the C5 concentration after two hours was 0.35 µmol/l with the UQ evaluation for the lipophilicity *l* and for the renal clearance *cl*_*urine*_. In contrast, the expected mean value of the C5 concentration after two hours was considerably lower with the UQ evaluation for 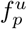 (0.25 µmol/l, Figure 4 (A)). Moreover, when comparing the C5 predictions of the same UQ evaluations in the urine, the urine clearance rate *cl*_*urine*_ showed the highest impact on the C5 prediction from the three evaluated parameters (Figure 4 (B)). The predicted C5 urine concentration varied between 0.006 - 0.012 µmol/l. The UQ evaluations with the fractional unbound plasma 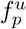 predicted urine C5 concentrations between 0.0058 - 0.01 µmol/l, whereas varying the lipophilicity *l* predicted a C5 concentration of 0.009 µmol/l in the urine (4 (B)).

**Figure 4:**
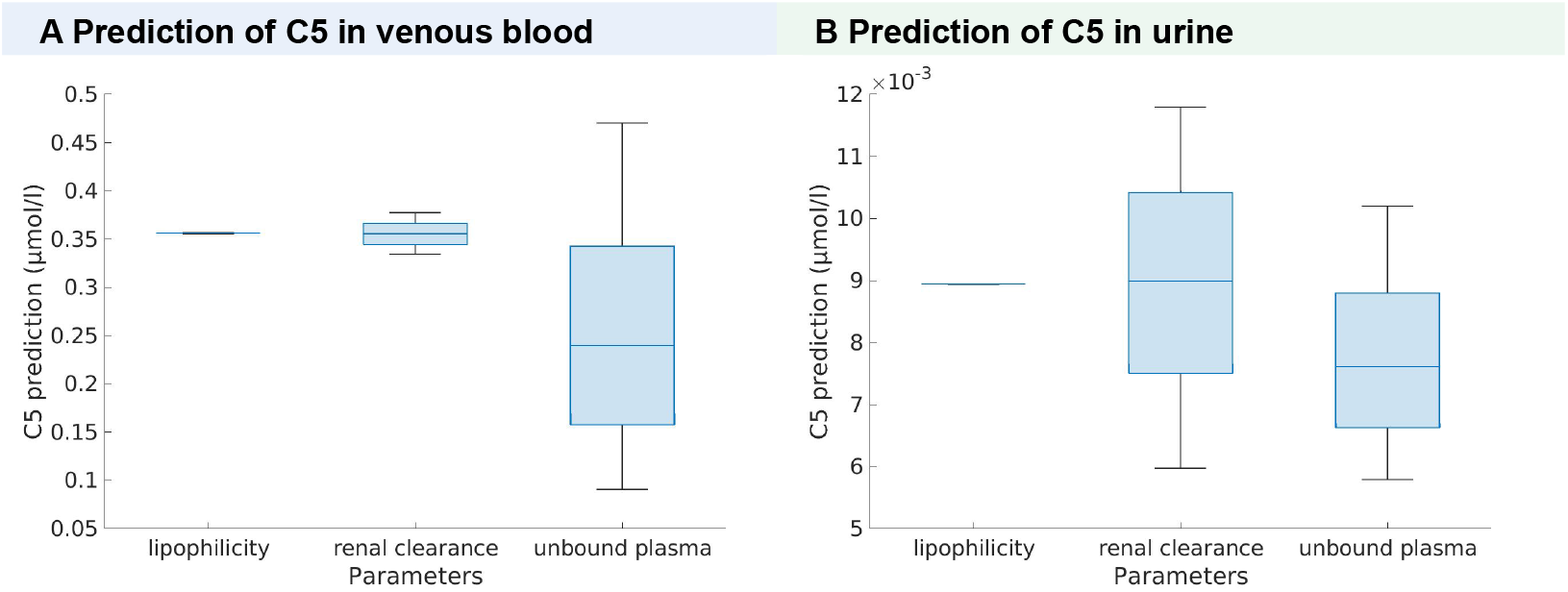
Boxplot of C5 prediction of PBPK model after 2 hours based on Monte Carlo Sampling for three uncertain parameters: lipophilicity, renal clearance rate, and unbound plasma rate in (A) the venous blood compartment and (B) the urine.

#### 3.4.5 Simulations with uncertain parameters

The UQ analysis revealed that the value of the fractional unbound plasma 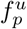 had a strong influence on the venous C5 prediction of the infant-specific PBPK model. Hence, we investigated what influence the choice of the fractional unbound plasma value would have on the results of the coupled dynamic infant-WBM model. For the previous evaluation the fractional unbound plasma was assumed to be 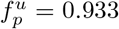. Therefore, we set the fractional unbound plasma to 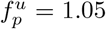, which was the upper bound of the sampling interval in the UQ analysis (section 3.4.3). This analysis on the WT and IVA model over a five hour interval as well as the prediction results on the 72-hour interval showed consistent trends as with the estimated parameter (Figure 5) although, as expected, the increase in C5 concentration was not as profound. The WT IMD simulation predicts a venous C5 concentrations after five hours of 0.3 µmol/l with 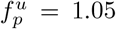. The IVA model shows a decreased ascent, the C5 prediction in the venous compartment after 5 hours is 0.45 µmol/l (Figure 5 (B)) whereas it was predicted as 0.59 µmol/l with 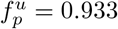 (Figure 2 (D)). After 72 hours, the predicted C5 concentration in the venous compartment is 1.02 µmol/l (Figure 5 (C)). Taken together, the choice of the fractional unbound plasma influences the IMD simulations for WT and IVA dynamic infant models but the overall result that the models predict increased C5 levels above the cutoff used by most newborn screening laboratories was not affected.

**Figure 5:**
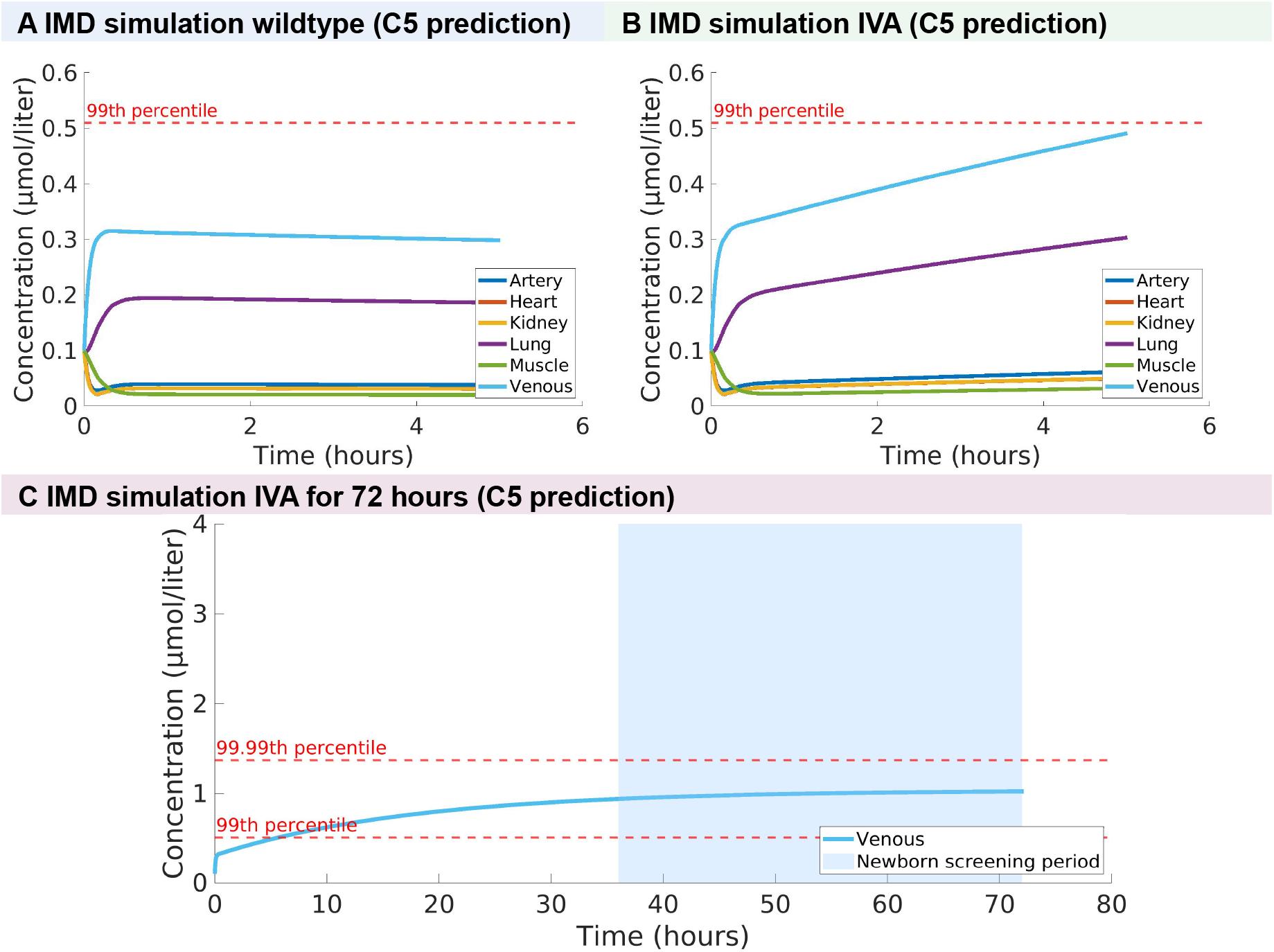
Concentration prediction of C5 from the dynamic infant-WBM for the artery and venous blood compartment, as well as the heart, kidney, lung, muscle over a five hour interval with the fractional unbound plasma parameter set to 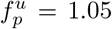. For an IMD analysis comparing (A) a wild type (WT) model and (B) an IVA model over five hours. And (C) an IVA model over a 72-hour interval also displaying the time interval, during which newborn screening is usually performed. The dashed lines show the 99^*th*^ percentile (0.51 µmol/l) and 99.99^*th*^ percentile (1.37 µmol/l) of blood C5 concentration used as cut-off values for IVA [53].

## 4 Discussion

We presented the development and application of dynamic infant-WBMs, which are an extension of the whole-body models for infantile metabolism [12]. The dynamic models incorporate PBPK modelling, which allows overcoming the steady-state assumption inherent to genome-scale metabolic models for selected compounds while expanding PBPK models for overall human metabolism. To illustrate the application of such dynamic infant-WBM, we developed a C5-specific dynamic model, which we used to predict changes in C5 bioavailability over time for the healthy and a disease state. We demonstrated that the dynamic model predicts increased blood concentrations for C5 in the disease state above the cut-off used in newborn screening and that this model could be used to predict dietary-based treatment strategies. As the dynamic infant-WBMs can be personalised based on an individuals parameters, they could enable personalised concentration predictions for known and novel biomarkers of healthy and diseased infant metabolism.

For the integration of time dependencies, an existing PBPK model [18] developed for drug research was utilised and adapted for infant physiology based on measured data [36] and adult references. In contrast to other dynamic metabolic models, which were based on a generic or cell-based model, such as Recon1 [29], the integration of a whole-body metabolic model allowed the mapping of organ-specific metabolite concentration predictions from the PBPK model to the constraint-based model. The developed procedure enables a flexible integration of different compartments, which can be selected depending on the investigated metabolite concentration, which allows for a compound-specific adaptation of the dynamic infant-WBM. We demonstrated this feature by developing a C5-specific dynamic infant-WBM that allowed for the prediction of the biomarker C5, including only the organs with C5 blood exchange reactions in the infant-WBMs. However, the reduction of the PBPK model to the overlapping organs and biofluid compartments could make model comparisons between different compounds more difficult, as the PBPK part of the model would be based on different ODEs. Hence, the compound-specific inclusion and exclusion of ODEs into the PBPK model should be evaluated carefully for every use case.

For the PBPK coupling, it was assumed that the predicted flux through the infant-WBM did not change every time step and was only updated every ten minutes for computational efficiency. When changing this evaluation interval to shorter intervals, the simulations showed no major differences in the C5 concentration prediction. However, this assumption could be inaccurate in scenarios where a rapid change of the metabolite flux is expected in a short time interval and should be reevaluated for different metabolites.

The dynamic infant-WBMs were parametrised with parameters calculated or obtained from experimental results in the literature. These parameters have uncertainties attached to them resulting, for example, from the experimental methods applied to obtain them. Hence, a UQ analysis for three of these parameters was performed to quantify the corresponding uncertainty. The Monte Carlo method highlighted the impact of variations of urinary clearance *cl*_*urine*_ and fractional unbound plasma 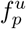 parameters on the predicted C5 concentration over two hours (Figure 3). The fractional unbound plasma 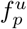 showed large variations in the blood C5 concentration with a mean of 0.25 µmol/l, which was considerably lower than the mean values (0.35 µmol/l) when the fractional unbound plasma was not varied. Consequently, it should be ensured that the fractional unbound plasma 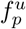 used in the dynamic infant-WBM is estimated correctly, as this parameter strongly impacts the output of the model. Similarly, the comparison between the C5 predictions in the urine compartment showed that variations in the urine clearance rate parameters have the strongest impact on the predicted concentration (Figure 3) and thus, this parameter should also be estimated correctly. Overall, the UQ analysis highlighted the importance of quantifying parameter uncertainties in mathematical models, such as the dynamic infant-WBMs, and that any conclusions derived from these simulations should consider the assumed parameters in the model.

The IMD research is evolving from a ‘static’ view of simple pathways towards a ‘dynamic’ view of metabolic fluxes [5]. Hence, we demonstrated how the dynamic infant-WBMs can enable a simulation of the dynamic behaviours of infant metabolism. We predicted the concentration changes of the clinically relevant biomarker C5 for isovaleric aciduria (IVA) in the healthy dynamic-infant WBMs, which reached steady state within the simulated two hour interval (Figure 2). Importantly, the predicted values (female: venous (0.38 µmol/l), lung (0.33 µmol/l; male model: venous: 0.31 µmol/l; lung: 0.19 µmol/l) remained below the 99^*th*^ percentile cut-off value (0.51 µmol/l) used in clinical practice for IVA [53] throughout the simulation time period. In contrast, in the dynamic infant-WBMs for IVA, in which we assumed no flux through the reactions associated with the genetic defect, the C5 concentration was seven times higher (i.e., 3.5 µmol/l, Figure 2) throughout the recommended age for newborn screening in Germany at 36 - 72 h of life [54]. However, these cutoff values represent only a guidance for further biochemical and genetic testing. In one large-scale study, individuals with attenuated IVA had a median C5 value of 2.6 µmol/l (range: 0.7 - 6.4 µmol/l) and individuals with classic IVA of 10.6 µmol/l (range: 4.3 - 21.7 µmol/l) [42]. In another study, the mean and standard deviation of C5 concentrations in newborns with attenuated IVA (n=22) was 2.6*±*1.16 µmol/l, whereas for severe patients with classic IVA it was 12.6*±*5.22 µmol/l [52]. A further study showed that C5 concentrations in dried blood spots above the cut-off of 5.6 µmol/l could be associated with a symptomatic classic IVA disease course with metabolic decompensations and an in trend inversely correlated IQ [43]. While the ranges of C5 in newborns with attenuated and classic IVA varied, they were all higher than our predictions after 72 hours. This highlights the difficulty of interpreting the dynamic infant-WBMs quantitatively. Moreover, as the UQ analysis showed, critical model parameters need to be either experimentally determined or multiple estimation approaches should be used to certify their values. These parameter uncertainties could possibly explain the discrepancy between the predicted and expected C5 concentrations. Nonetheless, our example demonstrated the potential use of the dynamic infant-WBMs for the qualitative prediction of C5 biomarker dynamics and how the developed procedure could be easily applied to other biomarkers and other IMDs associated with newborn screening.

The ability of infant-WBMs to predict the effect of dietary interventions on an IMD model has already been demonstrated [12]. This kind of treatment analysis needs to be patient-specific since the disease therapy and dietary guidelines for IMD patients have to be adapted to age-dependent requirements and the patient’s disease severity [5]. Here, personalised infant-WBMs can account for the substantial variability in patients since their treatment has to be individualised based on the patient’s diagnosis and clinical phenotype [5]. In this study, we showed that a reduction of dietary leucine decreased the C5 prediction in the IVA model. It should be noted that in our proof-of-principle, we could not decrease the dietary leucine intake below 10% of the normal infant diet applied as leucine is an essential amino acid and the model did not account for the infantile gut microbiota. In adult men, the microbial contribution to host leucine levels has been estimate to be between 24–27% of the estimated average requirement [61]. Expanding the dynamic infant-WBM to the gut microbial metabolic activity would be a valuable next step. The integration of personalised microbial community models with WBMs has been already demonstrated for the adult WBMs [11, 62, 63]. By progressing to dynamic infant-WBMs, and potentially even including gut microbial metabolism, would enable *in silico* predictions of short-term metabolic responses of an individual’s metabolism to the diet intake. By this, the models could support clinicians in personalised treatment planning and therapy for IMDs.

Moreover, the developed framework could be used for modelling pharmacokinetics and pharmacodynamics of drugs specifically for infants as these processes are often different from adults [64, 65]. This could allow for infant-specific drug dosage determination, which enables to account for the infant’s immature drug metabolism that is often associated with drug toxicity [66]. This is important since almost 50% of prescription drugs lack age-appropriate dosing guidelines [66]. Additionally, paediatric drug clearance data is less attainable, which is probably due to the difficulties associated with conducting paediatric clinical trials [67, 68]. Hence, analysing these drug-related metabolic processes *in silico* could be very beneficial for researchers, clinicians, and patients.

These results illustrate the great potential of dynamic infant-WBMs to predict temporal changes in known and novel biomarkers for IMDs. However, any *in silico* predictions need to be validated against experimental data and by clinical experts. In the case of the fractional unbound plasma, different modelling methods have been developed [69, 70, 71], which could provide more reliable values for this critical parameter. Furthermore, obtaining all the data necessary to personalise the dynamic infant-WBMs from one individual would allow for an improved personalisation of the dynamic infant-WBMs. Additional data, especially measurements from infants over several time intervals, are essential to enhance model validation and would contribute to the model’s reliability and applicability. In future studies, applying personalised data from patients with IMDs could be a next step to obtain quantitatively correct metabolite concentration prediction.

In conclusion, the dynamic infant-WBMs showcased the potential of coupling PBPK and infant-WBM models, opening new research directions for dynamic evaluations of infant metabolism. This capability holds significance for future infant-specific drug research and dietary treatment planning, where understanding the dynamic development of metabolite concentrations and the metabolic response to dietary interventions over time is crucial.

## 5 Methods

### 5.1 Physiological-based pharmacokinetic modelling

Physiological-based pharmacokinetic (PBPK) models extend Pharmacokinetic (PK) models, which are used to understand drugs and their distribution in an organism following an intravenous or oral dosing [35]. PK models describe the change of drug concentration *C* over time *t* depending on the drug clearance *Cl*, which describes the volume of plasma or blood that is cleared of the drug per unit time (l/min) and the volume of distribution *V*, which is the volume of the organ or tissue a drug distributes into [72],

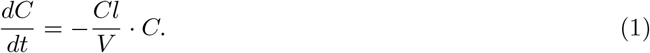

PBPK models enable the integration of various physiological parameters into this PK framework by adding compartments that correspond to different organs and tissues in the human body [35]. In the following, the term tissue *T* will be used for all organs and compartments of the human body considered in the PBPK models, such as organs or blood compartments. The tissues *T* are connected through the blood exchange in both venous and arterial blood. The arterial blood is the oxygenated blood, which is transported to all organs, except the lung (Figure 6). The venous blood is the deoxygenated blood, which leaves the organs and is pumped back into the lungs for oxygen uptake (Figure 6). For each tissue *T*, except for the lung, the rate of concentration change over time *t* of a compound *C*_*T*_ is described by the arterial inflow concentration *C*_*a*_ and the tissue-specific venous outflow concentration 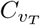,

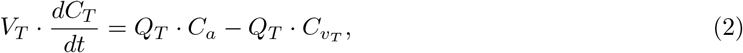

where *V*_*T*_ is the tissue volume (l), *Q*_*T*_ is the blood flow (1/h), and 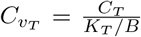 is the venous outflow concentration with *K*_*T*_ the tissue partition coefficient and *B* the blood-to-plasma ratio. The blood-to-plasma ratio is used to correct for blood when the plasma is used to determine the pharmacokinetic parameters, although the blood should be the true central compartment [73]. Based on this, the generic ordinary differential equation (ODE) of the rate concentration *C*_*T*_ in a tissue *T* that is changing over time *t* can be written as

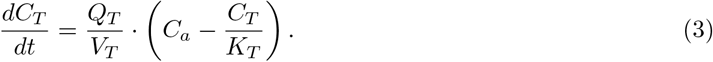

**Figure 6:**
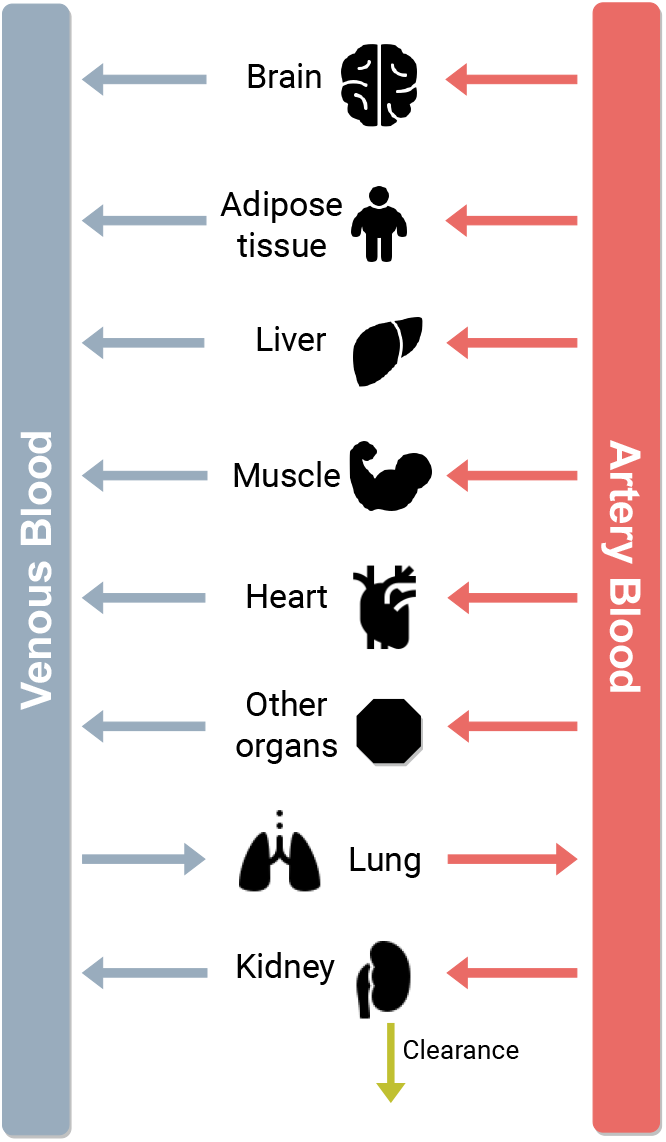
Overview of general PBPK model. Red arrows describe the blood exchange between artery blood and organs, blue arrows present the blood exchange between venous blood, and organs. The green arrow presents the renal and bile clearance.

Different approaches, such as perfusion rate-limited kinetics and permeability rate-limited kinetics, are used to model the kinetics within the compartments. In our work, all processes are modelled with perfusion rate-limited kinetics since it often occurs for small lipophilic molecules, which can dissolve easier in lipids than in water [35]. For these molecules, the blood flow to the tissue becomes the limiting process [35]. In perfusion rate-limited kinetics, it is assumed that tissue membranes do not impede diffusion. At steady state, the rate of drug entry into a compartment equals the product of the organ’s blood flow rate and the incoming blood concentration. The total drug concentration in a tissue *T* is in equilibrium with the concentration in the circulation as governed by the drug-specific tissue partition coefficient *K*_*T*_ [35]. The time to reach a steady state is determined by *K*_*T*_, the blood flow rate *Q*_*T*_, and the tissue volume *V*_*T*_ [35]. There are several methods to estimate the tissue partition coefficient *K*_*T*_. In this study, the Poulin method is applied [39], which assumes that the compound distributes homogeneously into the tissue and plasma by passive diffusion [35]. Based on the Poulin method [39], the tissue partition coefficient *K*_*T*_ of a tissue *T* can then be determined as,

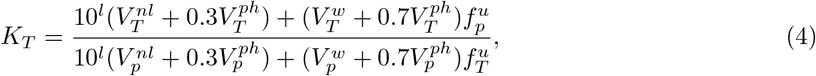

where *l* is the lipophilicity, referring to the capacity of a chemical compound to dissolve in lipids, 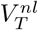 and 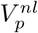 are the specific volume fractions of neutral lipid, 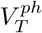 and 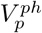 are the specific volume fractions of phospholipid, and 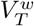 and 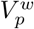 are the specific volume fractions of water in tissue *T* and plasma *p*, respectively [50]. All these volumes can be obtained from literature [50]. Furthermore, the chemical bind in plasma or fractional unbound plasma 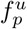 defines a drug’s binding degree in plasma [60]. This fraction describes the fraction of the compound that is not bound to plasma proteins and free for interaction with receptors, metabolizing enzymes, and renal filtration [73]. The compound-specific values for 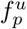 can be obtained from experimental results in literature [74] and applied to calculate the chemical bind in tissue, 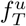, with

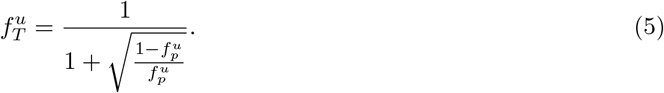

In addition to the tissue-specific models described by Eq. (3), the PBPK model incorporates additional equations for the arterial and venous blood compartment, as well as for the renal and bile excretion. The arterial and venous blood compartments connect all the tissue-specific ODEs (Figure (6)).

### 5.2 PBPK model used for the dynamic infant-WBMs

We used an existing PBPK model [18], which describes the dynamics of a system with 23 ODEs. For each compound of interest *i*, the time-dependent concentration in a specific tissue, such as adipose tissue, brain, small intestine, large intestine, heart, kidney, liver, lung, muscle, pancreas, skin, spleen, stomach, and bone, was calculated. Additionally, this PBPK model contained auxiliary ODEs to describe the transit from bile to small and large intestine lumen as well as separate equations for renal and bile clearance. For a compound *i* (a metabolite or drug), the applied PBPK model contains the following system of equations:

**Adipose tissue**

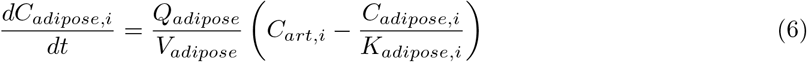

**Artery blood**

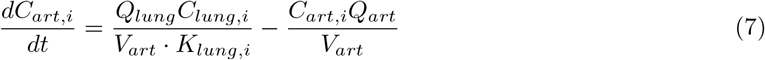

**Brain**

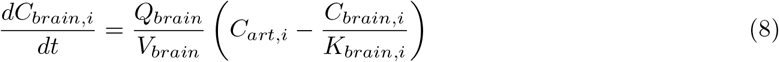

**Small intestine**

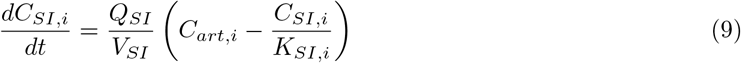

**Large Intestine**

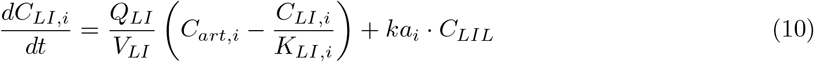

**Heart**

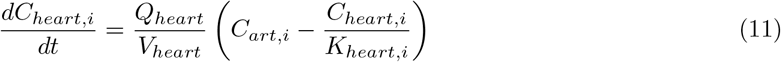

**Kidney**

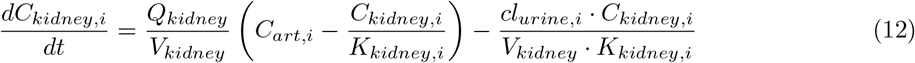

**Liver**

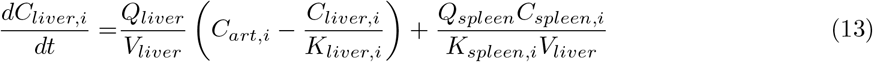

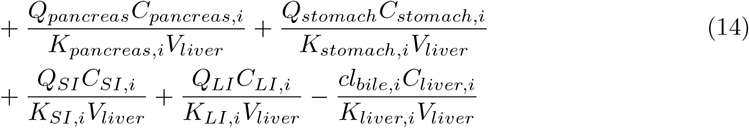

**Lung**

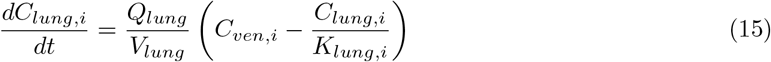

**Muscle**

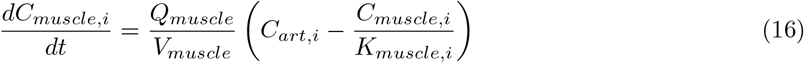

**Pancreas**

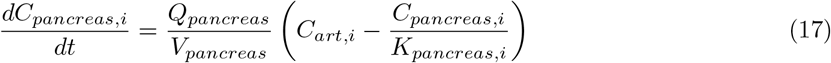

**Skin**

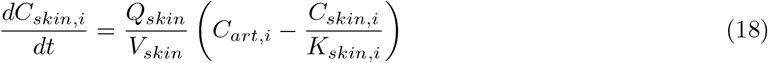

**Spleen**

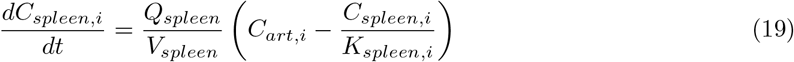

**Stomach**

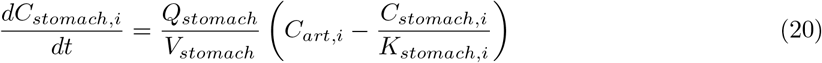

**Bone**

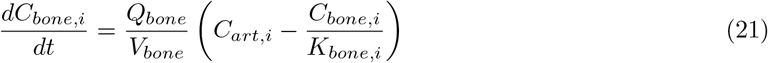

**Venous**

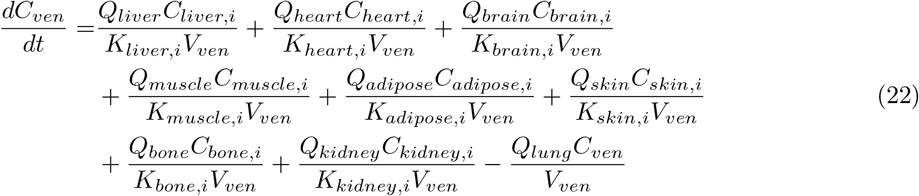

**Transit from bile to small intestine lumen**

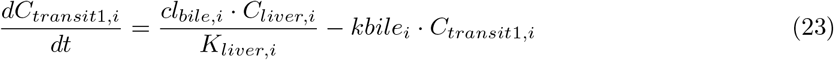

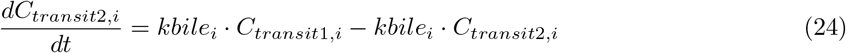

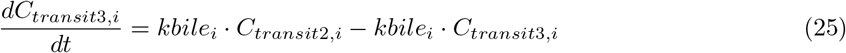

**Small intestine lumen**

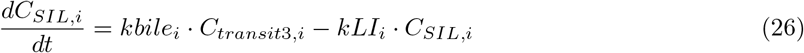

**Large intestine lumen**

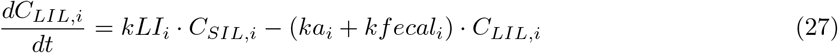

**Urine**

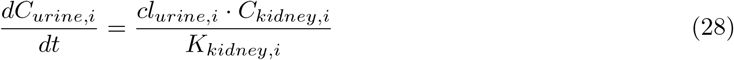

**Faeces**

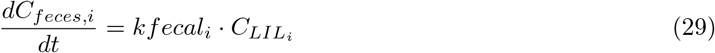

Here, *C*_*T,i*_ represents the concentration of the compound *i* in a specific tissue *T, V*_*T*_ is the tissue volume, *Q*_*T*_ the blood flow rate, and *K*_*T,i*_ the calculated organ-specific tissue partition coefficient (Eq. 4) for the compound *i*. Furthermore, *cl*_*urine,i*_ and *cl*_*bile,i*_ present renal and bile clearance for a specific compound, which are set to zero if the compound cannot be found in the urine or faeces, respectively. The compound-specific constants *ka*_*i*_, *kbile*_*i*_, *kfecal*_*i*_, and *kLI*_*i*_ are the large intestine, bile, and faeces coefficients, which are set to zero if the compound cannot be found in the faeces. This system of equations can then be solved by a commercial ode solver, such as the ode45s.m function in MATLAB [59].

### 5.3 Whole-body models for infants (infant-WBM)

The virtual metabolic human identifiers (VMH IDs) referred to in this study represent the reaction names of the corresponding organ-specific reactions in the infant-WBM models. The non-organ-specific part of the VMH ID can be searched in the VMH database (https://www.vmh.life) for further information. The metabolic whole-body models for infants (infant-WBM) [12] present a resource of sex-specific, organ-resolved whole-body models of infant metabolism spanning the first 180 days of life. For the creation of these models detailed knowledge of infants’ physiology, metabolic processes, and newborn screening data was used. They account for organ-specific parameters and included in detail the energy demand for brain development, heart function, muscular activity, and thermoregulation. The models represent active infant metabolism and predict infant growth between zero and six months in agreement with growth trajectories by the WHO [13]. In this study, a male and female infant-WBM at age one day were used from the metabolic whole-body models for infants (infant-WBM), resource (https://www.vmh.life/files/Newborns/) [12]. The male infant-WBM has a weight of 3.3 kg, a predicted growth rate at day one of 0.0089%, and was fed 122 ml of breast milk on that day. The female infant-WBM has a weight of 3.2 kg, a predicted growth rate of 0.0075%, and was also fed 122 ml of breast milk. All physiological and nutritional parameters of both models are presented in Table S4.

#### 5.3.1 Flux balance analysis

For the flux balance analysis (FBA), the hypergraph of a metabolic network is converted into a stoichio-metric matrix *S* ∈ ℝ^*m*×*n*^, where the rows correspond to the *m* metabolites and the columns to the *n* reactions. Each matrix entry *S*_*ij*_ corresponds to a stoichiometric coefficient, which indicates if and how metabolite *i* takes part in reaction *j*. The change of a metabolite concentration *x*_*k*_ over time *t* is then represented by the *k*^*th*^ row, while the computed flux value *v*_*i*_ corresponds to the *i*^*th*^ entry in the flux vector *v* = (*v*_1_, …, *v*_*n*_)^*T*^. Resulting in the mass-balance equation for all metabolites as

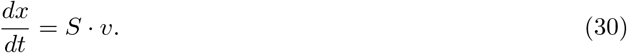

In this study, we use the COBRA approach [14], which assumes the simulated system to be at a steady state and, hence,

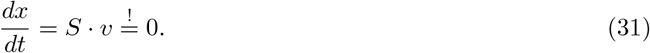

This linear system of equations is then used as constraints in flux balance analysis (FBA) [15], which is a mathematical approach to minimise or maximise an objective function, e.g., biomass yield, through the metabolic network defining the following linear program (LP),

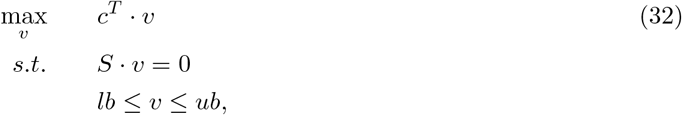

where *lb* and *ub* are the lower and upper bounds on the flux vector *v*. These bounds are based on specific data, such as physiochemical constraints, nutrient uptake rates, and enzyme reaction rates. The vector *c* is a vector of weights indicating which and how much each reaction *i* contributes to the objective function [15]. By definition, the lower bound on an irreversible reaction was set to *lb* = 0 mmol/day/person and *ub >* 0 mmol/day/person [15, 11]. For reversible reactions, negative flux through the reactions was allowed, *lb <* 0 mmol/day/person and the upper bound was set to *ub >* 0 mmol/day/person. The *lb* and *ub* of unconstrained reactions were set to the arbitrary values -1,000,000 mmol/day/person and 1,000,000 mmol/day/person, respectively.

### 5.4 Four-step procedure for coupling PBPK models and infant-WBMs

For the coupling of PBPK models and infant-WBMs, we developed a four-step procedure that can be applied to any metabolite biomarker of interest, such as C5. Overall, the flux through a metabolic reaction in the infant-WBM was interpreted as the intrinsic ability of a compound to be metabolised by the relevant enzyme. Hence, the flux was used to update the PBPK model for predicting the metabolite concentration (Figure 1). Likewise, the predicted metabolite concentrations were used to update the bounds on the flux values in the infant-WBM. This led to an iterative scheme, where both models were evaluated one after the other, and the predictions were used to update the models. This iterative scheme allows to combine information from physiology, pharmacokinetics, and metabolism. The following four steps describe the coupling of both models leading to time-dependent infant-WBMs.

#### 5.4.1 Step 1. PBPK adaptations for infant physiology

First, a PBPK model [44] is adapted for specific physiological parameters corresponding to an individual. These parameters are extracted from the individual parameters variable of the infant-WBM. Similar to Toroghi et al [29], the physiological and physicochemical properties in the PBPK model are assumed to be constant over time. In particular, the PBPK parameters for the body mass, organ volumes, and organ flow rates are obtained directly from the infant-WBM models. The calculation of the organ volumes is based on the organ weights reported in the infant-WBM and the respective organ density obtained from literature [36]. The venous and arterial blood volume estimations are based on the total blood volume resulting in 144 ml of blood in both compartments for a 3.3 kg newborn at day 1 (total blood volume: 288 ml). This is comparable to the blood range of newborns at 72 hours of age (75 to 107 ml/kg) ranging between 248 - 353 ml for a 3.3 kg infant [47].

The organ flow rates are obtained from the organ flow rate parameters of the infant-WBM, which are measured in l/min/kg tissue and converted to l/h/organ. The blood flow rate for venous and arterial blood is estimated as 3 l/h since the blood flow rate ranges from 10 to 50 ml/min in infants weighing less than 5 kg [38], leading to an estimation of 0.6 - 3 l/h by multiplication with 60/1000. The body surface area is calculated based on the Mosteller method [48], using the average height of 50 cm for male and 49 cm for female newborns [37].

#### 5.4.2 Step 2. Defining compound-specific parameters

In the second step, the compound-specific parameters are set. A compound of interest can be a drug or a biomarker, such as an amino acid or acylcarnitine. All processes are modelled with perfusion rate-limited kinetics, and the Poulin method [39] is used to calculate the tissue partition coefficient *K*_*t*_, section 5.1. For every compound, the molecular weight, lipophilicity *l*, and fractional unbound plasma 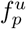 have to be determined. The molecular weight and lipophilicity *l* are obtained from databases, such as HMDB [40]. The fractional unbound plasma 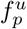 can be measured experimentally for amino acids [74] and acylcarnitines [49] evaluating the recovery rate, which is obtained by comparing results before and after spiking with known concentrations of standards in plasma [49]. Furthermore, the renal and bile clearance rate (l/h) is extracted from the literature for amino acids [75] and L-carnitine [41]. To account for the consumption and production rate of a compound in an organ, *R*_*organ*_, which is characterized by the Michaelis-Menten kinetics,

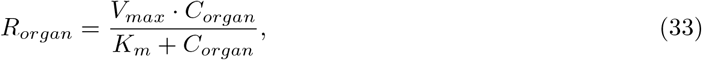

is subtracted in each organ of the PBPK model [29, 24]. It is defined by, the compound concentration *C*_*organ*_ in a specific organ, *V*_*max*_ the maximum velocity of the compound-producing reaction, when all the enzyme’s active sites are saturated with substrate, and *K*_*m*_ the Michaelis-Menten constant for the compound [24].

#### 5.4.3 Step 3. Time-dependent model parameters

To integrate time dependency into the models, the length of the total time intervals and the time intervals calculated by the ODE solver have to be defined. Furthermore, the frequency of the update of the infant-WBM to the ODE model has to be determined. The ODE system of equations can be solved with the ode45s.m function in MATLAB [59, 76].

#### 5.4.4 Step 4. Iterative integration of PBPK and infant-WBM

Steps 1. - 3. assemble an infant-specific PBPK model for a specific compound of interest. To integrate the infant-WBM and the PBPK model, the latter is iteratively updated using flux predictions from the infant-WBM, which in turn is revised based on the predicted metabolite concentrations from the PBPK model. This integration is detailed in the following sub-steps.

##### Step 4.1 Set initial values for compound concentrations in organs

The initial value of the compound concentration of interest is set to a published blood concentration measurement of this compound. This value is set for the concentration in every organ and compartment that has an exchange reaction from the organ to the blood compartment (VMH ID: ‘EX *compound* [bc]’). Additionally, the concentration of the artery and venous blood compartment is set to the initial concentration. The ODEs representing the renal and bile clearance are added to the PBPK model depending on the excretion of the compound of interest. All other tissue-specific ODEs from the full PBPK model are excluded from the compound-specific PBPK model.

##### Step 4.2 Compute metabolite concentration with PBPK model

The PBPK model computes the metabolite concentration for a set time frame. Hence, a dynamic concentration prediction on the PBPK model assembled in steps 1-3 is solved with the ode45s solver implemented in MATLAB [76, 59].

##### Step 4.3 Update bounds of infant-WBM with PBPK prediction

After the ODE time interval, the predicted metabolite concentration (in mmol/l) is utilised to update the infant-WBM bounds. Therefore, the predicted blood concentrations *C*_*art*_, *C*_*ven*_ are used to set the lower bounds on the uptake flux *lb*_*organ*_ of the compound from the blood circulation into individual organs utilising the organ-specific plasma flow rate (*PFR*_*organ*_).

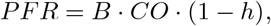

where *B* is the organ-specific blood flow rate, *CO* is the cardiac output and *h* is the hematocrit value. For the lung, the venous compound concentration is used since it transports blood from the organs to the lung

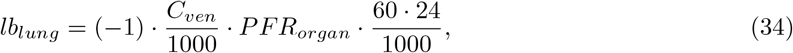

given in mmol/day/person. For all other organs, the artery compound concentration is applied since the arterial blood is the oxygenated blood transporting oxygen from the lungs to the organs.

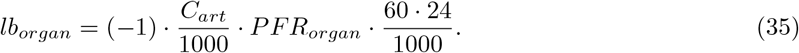

By restricting the lower bound of the reactions, the uptake of the biomarker from the respective blood compartment into the organ is restricted.

##### Step 4.4 Run infant-WBM LP optimisation

After step 4.3, the infant-WBM is updated based on the PBPK model and can be evaluated. The whole-body biomass reaction is set to the maximal growth rate of the infant on the day. Then, the LP for the infant-WBM is solved, minimizing the fluxes while adhering to all constraints to obtain a flux *v*_*j*_ for each reaction *j* in the network.

##### Step 4.5 Update PBPK model with infant-WBM flux prediction

From the LP solution, the predicted flux *v* through the exchange reaction of the compound are extracted. First, these fluxes are computed in the unit mmol/person/day and then converted to mmol/volume/second. Second, the resulting value is divided by the respective organ volume, as the concentration change over time in the PBPK model is given per organ volume. By this, the converted flux 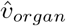 for each organ and the respective excretion pathway is obtained which is used to update the PBPK model. Therefore, the flux is subtracted in the ODE representing the compound in each organ,

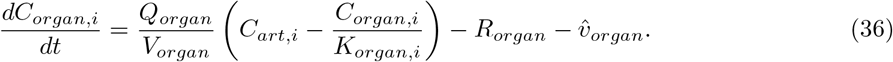

Then, the evaluation starts again at step 4.1.

### 5.5 IMD analysis for dynamic infant-WBMs

The IMD analysis is utilised in metabolic modelling to demonstrate the human metabolic model’s capability to predict known biomarkers. IMDs often disrupt the normal metabolite flux. Hence, analysing the metabolic flux over time is more informative for IMDs than the static measurement of metabolites at a single time point [5]. In this work, we used the IMD analysis method for whole-body models, established in Thiele et al [11], which is a two-step process. In the first step, all reactions *k* = (*k*_1_, …, *k*_*n*_)^*T*^ in all model organs associated with the defect gene that causes the IMD are identified. The contribution of these reactions *k*_*i*_ is then summed in an artificial reaction *v*_*d*_. This reaction is added to the model, and a first LP is solved, maximizing *v*_*d*_. In the second step, 75% of the optimal solution *z*^∗^ is used as a lower bound on *v*_*d*_ in the wild type (WT) model. And in the IMD model the flux *v*_*k*_ through the *k* associated disease reactions is set to zero. Second, for both models, an artificial demand reaction in the blood compartment *v*_*D*_ is set as objective function for flux maximization. The LP of the WT model can be written as,

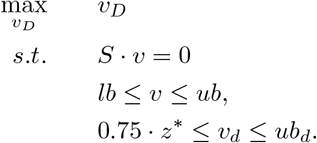

This LP then obtains the WT optimal flux through the blood demand reaction 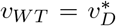. The IMD model can be summarised as,

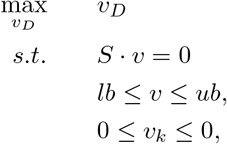

where 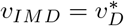 is set to the optimal flux value.

For the dynamic IMD analysis, this IMD simulation is performed in every FBA step and the solution is used to update the PBPK model iteratively. Therefore, the optimal fluxes *v* _*WT*_ and *v*_*IMD*_ are added to the venous blood compartment with a correction factor *c* = 0.1, which relaxes the maximisation of the blood demand of a specific compound. Hence, the ODE of the venous blood compartment f or the IMD analysis of the WT model can be given as,

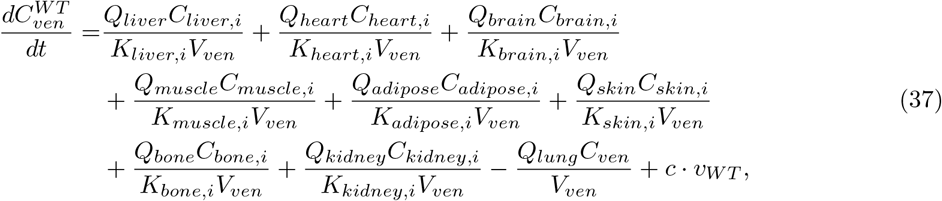

and for the IMD model as

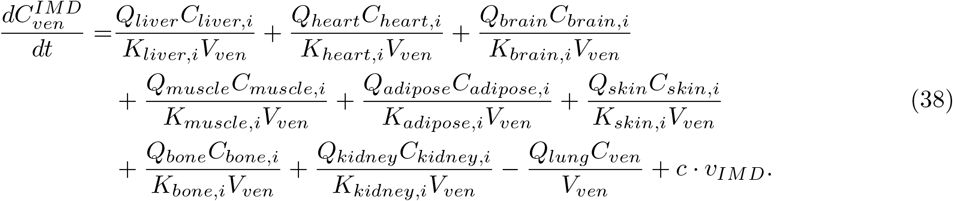

### 5.6 Uncertainty Quantification

Mathematical models, such as infant-WBMs and PBPK models, attempt to simulate complex biological processes. These models are developed using necessary assumptions about process behaviours, parameters from the literature, and parameter approximations. However, these assumptions and simplifications introduce uncertainties into the models. Uncertainty quantification (UQ) analyses all sources of error and uncertainty in mathematical models. This process assesses the extent to which a model should be trusted rather than determining whether the model is correct or incorrect [77]. UQ can help assess an uncertain parameter’s impact on a model by studying the relationship between the uncertainty in the output and input of a model [78].

#### 5.6.1 Monte-Carlo Simulations

To analyse parametric uncertainties in a mathematical model such as the dynamic infant-WBMs, computational methods, such as Monte Carlo simulation, can be applied. This non-intrusive method propagates uncertainties through a mathematical model [58]. It is based on random sampling. An underlying probability distribution with a probability density function *f* is chosen for this. This distribution could be a continuous uniform distribution in an interval *I* = [*a, b*],

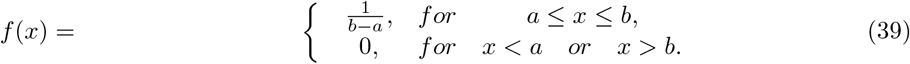

Such a uniform distribution can be applied when there is a minimum and maximum value in which the parameter arbitrarily lies. Then, *N* input random variables *Z* ∈ ℝ^*d*^ are generated according to the chosen probability density function *f*. The parameters *Z*^(*i*)^ with *i* = 1, …, *N* are used as input to the model *m*(*Z*^(*i*)^). When evaluating the model output *m*(*Z*^(*i*)^), the impact of the variation of the parameter in the interval *I* on the model output can be assessed. Using these model realisations, the quantity of interest, such as the expected value, can be computed, [58]

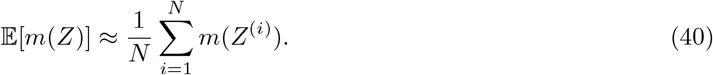

## Acknowledgements

The authors acknowledge the support of the Informatics for Life project funded by the Klaus Tschira Foundation, the financial support by the Dietmar Hopp Foundation, St. Leon Rot, Germany (grant numbers 2311221, 1DH2011117, and 1DH1911376), the financial support by the European Research Council (ERC) under the European Union’s Horizon 2020 research and innovation programme (#757922 and #101125633), and the Horizon Europe grant Recon4IMD (#101080997). The present work was also supported by the Helmholtz Association under the joint research school HIDSS4Health – Helmholtz Information and Data Science School for Health.

## Author contribution

Conceptualisation E.Z., F.K.M., V.H., I.T.; Methodology E.Z., F.K.M., V.H., I.T.; Software E.Z., F.K.M., I.T.; Data Curation E.Z., F.K.M., I.T.; Writing – Original Draft E.Z., I.T.; Writing – Review & Editing E.Z., F.K.M., U.M., S.K., V.H., I.T.; Supervision U.M., S.K., V.H., I.T.; Funding Acquisition U.M., S.K., V.H., I.T.

